# Instant Taq: Rapid Autoinducible Expression and Chromatography-free Purification of Taq polymerase

**DOI:** 10.1101/2021.09.25.461774

**Authors:** Romel Menacho-Melgar, Tian Yang, Michael D. Lynch

**Affiliations:** Department of Biomedical Engineering, Duke University

**Keywords:** Taq, PCR, Autolysis, Autohydrolysis, Purification, Chromatography-free

## Abstract

DNA modifying enzymes are ubiquitous reagents in synthetic biology. Producing these enzymes often requires large culture volumes, purified nucleases and chromatographic separations to make enzymes of necessary quality. We sought to leverage synthetic biology tools to develop engineered strains allowing for not only the production but rapid purification of these reagents. Toward this goal, we report an *E. coli* strain enabling the rapid production and purification of Taq polymerase. The method relies on 1) autoinducible expression achieving high protein titers, 2) autolysis and auto DNA/RNA hydrolysis via lysozyme and a mutant benzonase™, and 3) heat denaturation under reducing conditions to precipitate contaminating proteins including the mutant benzonase™. Taq polymerase is obtained at high purities (>95% pure by SDS-PAGE) and is readily usable in standard reactions. The method takes less than 1 hour of hands-on time, does not require special equipment, expensive reagents or affinity purification. We expect this simple methodology and approach will improve access not only to Taq polymerase but to numerous additional commonly utilized reagent proteins.

**Highlights:**

- Protein titers ~ 1g/L achieved in 20 mL shakeflasks.
- 4 mg of purified Taq (corresponding to 5,000 Units, or 4,000 PCR reactions) per 20 mL shake flask.
- Instant Taq is equivalent to commercial preparations in routine PCR

## Introduction

Research enzymes are required for numerous methodologies ranging from DNA amplification to routine cloning.^1^ These reagents are indispensable in modern biology. Taq polymerase and other thermostable polymerases are perhaps the most ubiquitous and readily available research enzymes, due in part to numerous advances in expression and purification that have been made over the past 3 decades since Taq was cloned.^2,3^ Despite the advances in production and accessibility, even bulk Taq costs are still >$0.05/Unit with more pricey products such as Master Mixes costing as much as >$0.25/Unit. For labs performing large numbers of PCR reactions or other pertinent workflows, these costs are non-trivial. In our lab alone, the annual expenditure of Taq and higher fidelity polymerases can easily exceed $5,000-$10,000. When considering all reagent enzymes beyond polymerases, the costs can be much greater. While these reagents are ubiquitous, their costs still limit routine workflows in many labs as well as large scale applications even in the most well funded labs. More broadly, improving the accessibility of these core reagents will aid in the democratization of synthetic biology.

Although numerous methods have been reported for the in house expression and purification of Taq, they are not practical or cost effective for many labs, requiring significant time (often more than a day of hands-on time), large culture volumes (from 1 to 12 liter cultures), and often specialized reagents and/or equipment for cell lysis and chromatography, for example.^4,5^ In the case of Taq polymerase, one of the easiest methods, a recent ethanol based precipitation, still requires large culture volumes, timed induction and expensive reagents including purified lysozyme, DNAse and RNAse to lyse cells and remove contaminating nucleotides, which is essential for downstream PCR.^6^ Even trace amounts of contaminating bacterial DNA are known to complicate applications such as diagnostics or evolutionary studies.^6–8^

In this study, we sought to leverage advances in synthetic biology in combination with standard optimization approaches to develop an easy, readily accessible method for the routine in-house production of high quality polymerases. Toward this goal, we built upon strains and processes we have recently reported for the 1) autoinduction of heterologous protein expression upon phosphate depletion,^9,10^ 2) the simultaneous induction of a lysozyme and a broad specificity endonuclease (*S. marcescens nucA*, benzonase™) which enables the autolysis and autohydrolysis of host DNA/RNA,^11^ and 3) the simultaneous heat denaturation and precipitation of contaminants enabling purification of thermostable proteins.^12^ A combination of these approaches can be used to rapidly purify thermostable reagent enzymes as illustrated in Figure 1. While the nuclease is necessary in this approach to eliminate host contaminating oligonucleotides, removal of nucleases in the final purified enzyme preparation is critical in any downstream application of DNA modifying enzymes. A key advance in this work is the use of a mutant monomeric benzonase, with decreased thermostability, in combination with a reducing agent to enhance its precipitation, after the host DNA and oligonucleotides have been enzymatically removed.^13^ Together these advances enable both auto DNA/RNA hydrolysis during lysis and subsequent removal of contaminating endonuclease activity by heat inactivation/precipitation (Figure 1b).

**Figure 1.**
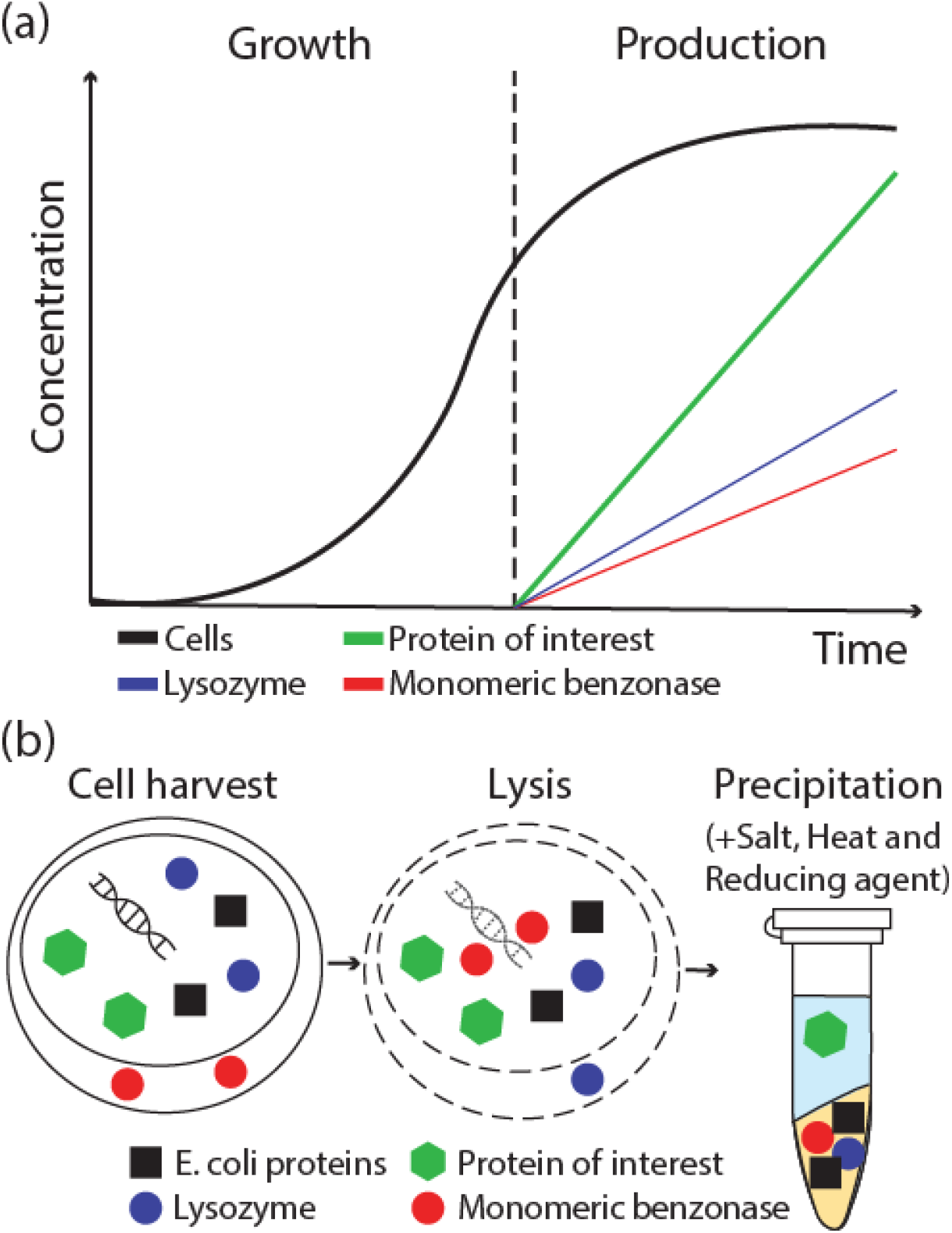
Rapid Autoinducible Expression and Chromatography-free Purification of thermostable proteins. (a) Two-stage expression of a protein of interest where in a first stage (“Growth”) the cells are grown until phosphate depletion in the media triggers the autoinducible expression of the protein of interest (second stage, “Production”), a lysozyme and a mutant monomeric benzonase. (b) After cell harvest, the cells can be autolysed and the host oligonucleotides removed leveraging the lysozyme and benzonase. After lysis, addition of salt, a reducing agent and heat causes the precipitation of all contaminants leaving a preparation of a purified thermostable protein of interest.

We have validated this approach via the production and purification of Taq polymerase, widely used in molecular and synthetic biology. The methodology results in high titers, yield and purity of Taq polymerase readily usable in downstream applications. In many respects, as simple as a routine plasmid DNA miniprep, yet requires no special columns or affinity reagents; only an incubator shaker, a heat block, a freezer and a centrifuge, all of which are basic molecular biology lab equipment.

## Results

We began by initially optimizing the process for Taq DNA polymerase, as illustrated in Figure 1. Firstly, to express Taq, we leveraged strains, expression constructs and protocols we have recently reported for the tightly controlled, low phosphate autoinduction of heterologous protein, as well as autolysis and autohydrolysis of DNA/RNA.^9–11,14^ Specifically, we cloned Taq into an expression construct driven by the robust, low phosphate inducible *E. coli yibD* gene promoter. ^9,15^ This plasmid was transformed into strain DLF_R004 (Genotype: F-, λ-, Δ*(araD-araB)567, lacZ4787(del)(::rrnB-3), rph-1, Δ(rhaD-rhaB)568, hsdR51, ΔackA-pta, ΔpoxB, ΔpflB, ΔldhA, ΔadhE*, *ΔiclR, ΔarcA, ΔompT::yibDp-ƛR-nucA-apmR*), which has been engineered for the low phosphate induction of a periplasmic nuclease (*S. marcescens nucA, benzonase™*) and cytoplasmic Lambda phage lysozyme as illustrated in Figure 1a.^11^ After the disruption of the membrane in the presence of 0.1% triton-X, lysozyme degrades the peptidoglycan cell wall and the nuclease degrades residual cellular DNA and RNA (Figure 1b).^11^ Cell growth and autoinduction of Taq expression is performed in autoinduction broth as described by Menacho-Melgar et al.^10,14^ Using this system, Taq was expressed to ~35% of the total cellular protein approaching titers of 900 mg/L as seen in Figure 2a.

**Figure 2.**
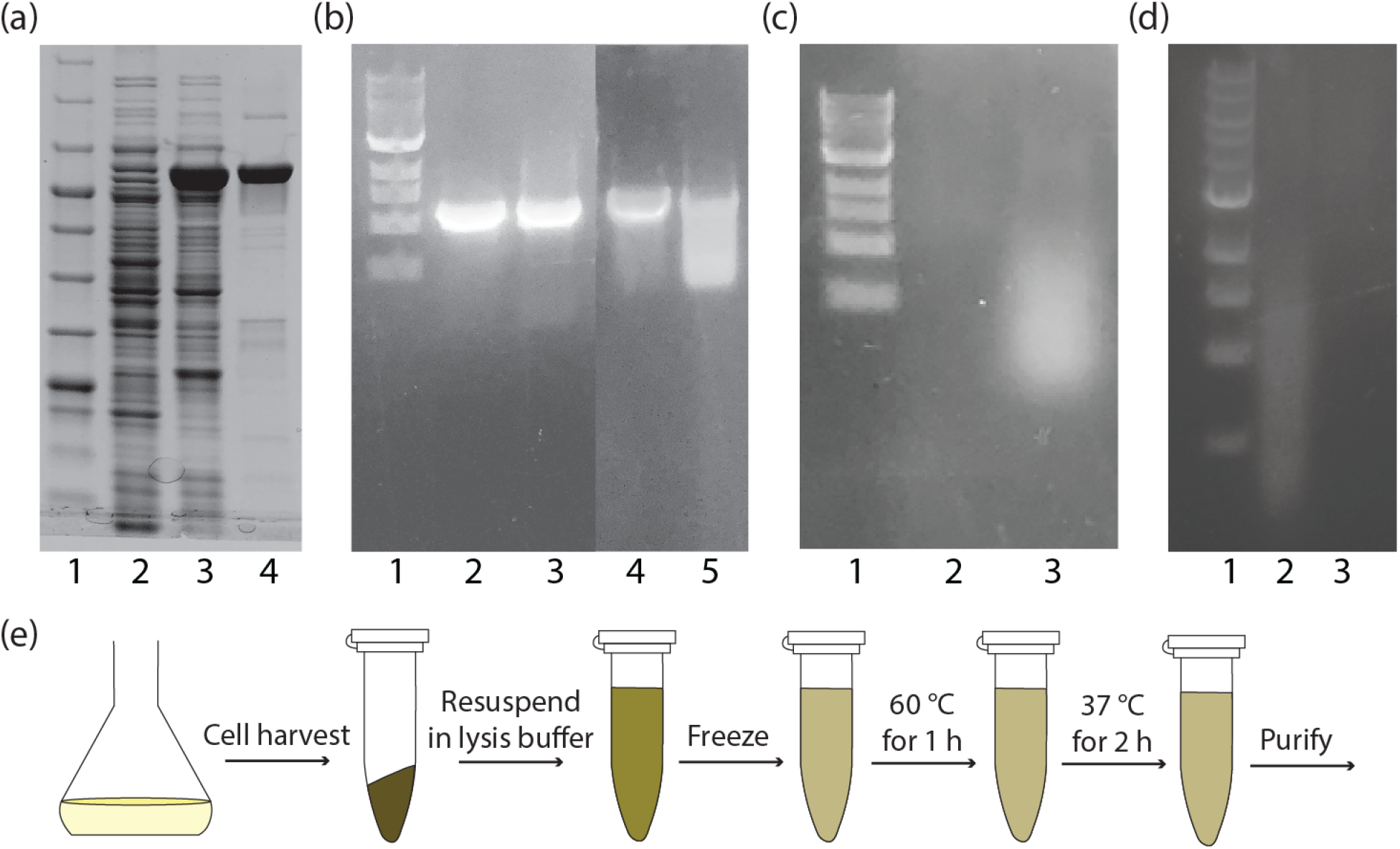
Residual nuclease activity and contaminating DNA in initial Taq preparations. a) Taq expression and initial purification using *E. coli* strain DLF_R004 (lane 1, ladder; lane 2, empty vector control; lane 3, Taq expression; lane 4, Heat extract after 30 minutes at 70 °C). b) Degradation of PCR product made using purified Taq initial preparations in comparison to commercial preparations. A PCR product made using commercial Taq (2) and initial Taq purifications (3), immediately after the PCR reaction and a PCR product made using commercial Taq (4) and initial Taq purifications (5) after incubating the PCR reaction overnight at 4 °C. c) Expression of Taq in DLF_R004 leads to contaminating DNA (lane 3) compared to an empty vector control (lane 2). d) Addition of a 60 °C step in the autolysis/autohydrolysis protocol (lane 3) leads to complete DNA removal when compared to the regular, non-modified autolysis/autohydrolysis protocol (lane 2). e) Schematic of the modified autolysis/autohydrolysis protocol where a 60 °C step is added after the freezing step, before incubating the cells at 37 °C

Despite good expression, an initial test of DNA/RNA autohydrolysis in strain DLF_R004 expressing Taq resulted in residual contaminating DNA in the lysate as shown in Figure 2c. We hypothesize that this contaminating DNA may be the result of benzonase producing short oligonucleotides, which can anneal and be extended by Taq at 37 °C. Thus, we modified the previously reported autolysis protocol ^11^ and included an incubation step at 60 °C for one hour after freezing which led to complete DNA/RNA autohydrolysis (Figure 2d). A schematic showing the modified autolysis/autohydrolysis protocol is shown in Figure 2e.

With successful autolysis and auto DNA/RNA autohydrolysis, we next turned to the further purification of Taq. Initial attempts at Taq purification relied on simply heating lysates to 70°C for 30 minutes followed by centrifugation to denature and precipitate contaminating proteins.^4^ Unfortunately, these original preparations contained residual contaminating non-specific thermostable endonuclease (*S. marcescens nucA)* activity, leading to the degradation of PCR product over time (Figure 2b). This contaminating activity is of course not compatible with downstream applications, including, in the case of Taq, routine PCR. In previous reports the non-specific DNA/RNA endonuclease from *S. marcescens*, requires very harsh conditions for complete inactivation (heat denaturation at 70°C for 30 min in the presence of 0.1 M NaOH).^16^ Elevated temperature as well as the addition of sodium hydroxide to our heat inactivation step were able significantly reduce nuclease activity albeit at a complete loss of Taq activity. (Figure S1). As a consequence we evaluated several options to remove contaminating nuclease activity. These included Taq purification using diafiltration with different membrane MW cutoffs and aqueous two-phase extraction. Unfortunately, these methods were unable to remove residual nuclease activity and DNA degradation (Figure S2-3).

Lastly, as benzonase has crucial disulfide bonds required for correct folding and activity^17^ (Figure 3a), we investigated the potential of adding reducing agents during the heat denaturation step to facilitate inactivation and precipitation of residual benzonase. The addition of 0.5M beta-mercaptoethanol during heat denaturation greatly improved, but still did not eliminate, contaminating endonuclease removal, as shown in Figure 3b. However, based on this improvement, we constructed a new host strain, DLF_R005 (Genotype: F-, λ-, Δ*(araD-araB)567, lacZ4787(del)(::rrnB-3), rph-1, Δ(rhaD-rhaB)568, hsdR51, ΔackA-pta, ΔpoxB, ΔpflB, ΔldhA, ΔadhE*, *ΔiclR, ΔarcA, ΔompT::yibDp-ƛR-nucA-apmR*), identical to DLF_R004, wherein the wildtype dimeric benzonase was replaced with a monomeric variant having H184R mutation.^13^ This variant is not only active as a monomer but has a lower melting temperature, making it a better candidate for thermal inactivation under reducing conditions. We confirmed the ability of this strain to undergo autolysis/auto DNA/RNA hydrolysis as well as enable Taq autoinduction (Figure S4). Importantly, strain DLF_R005 also enabled further reductions in nuclease activity (Figure 3b).

**Figure 3.**
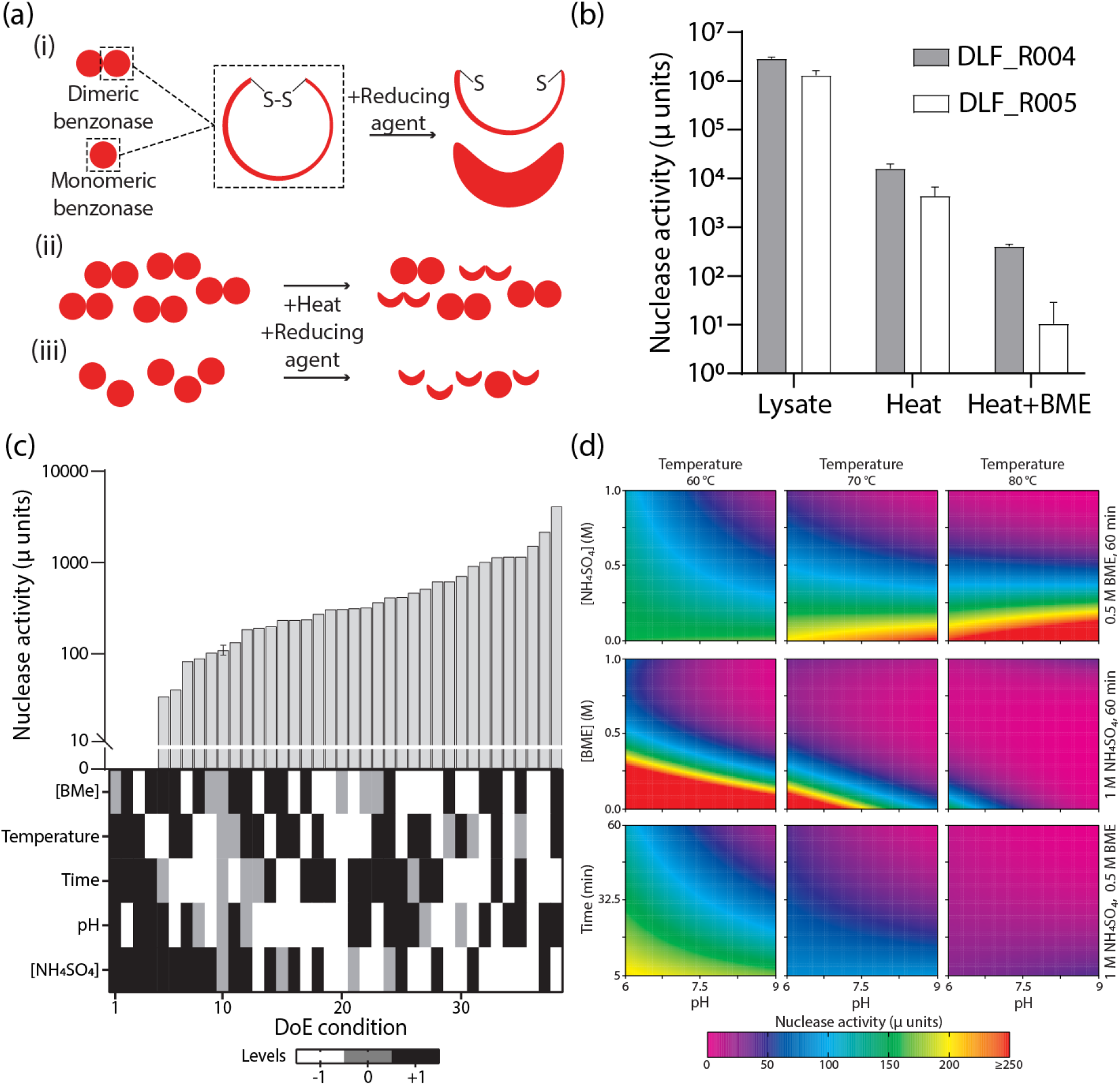
Optimization of nuclease removal. (a) (i) Essential to activity disulfide bonds in both dimeric (wild-type) and monomeric (mutant) benzonase are susceptible to a reducing agent causing loss of enzymatic activity. (ii) The combination of heat with reducing agent results in a permanent loss of enzymatic activity for the wild-type benzonase, but (iii) even lower residual activity for the mutant monomeric benzonase. (b) Quantification of residual nuclease activity (micro units) in lysates prepared from either strain DLF_R004 (dimeric benzonase) or DLF_R005 (monomeric benzonase) including: untreated lysates, lysates after 20 minutes at 60 °C, and heat treated lysates in the presence of 0.5 M beta-mercaptoethanol (BME). (c) Optimization of nuclease removal using Design of Experiments (DoE) using 38 different precipitation conditions, each comprising different ‘Levels’ of beta-mercaptoethanol (0 to 1 M), temperature (60 to 80 °C), time (5 to 60 minutes), pH (6.0 to 9.0) and ammonium sulfate (0 to 1 M). The upper panel shows nuclease activity ranked from lowest to highest activity. The lower panel shows the precipitation condition for each result. Refer to Table S3 for activity values and specific numbers for each Level within each experiment. All measurements were n=1, except for the centerpoint which was repeated 3 times to estimate experimental error. (d) Model based on DoE results. For all 9 plots, pH is varied across the x-axis from 6 to 9. For all plots within the same column, the temperature is the same and fixed at 60 °C, 70 °C or 80 °C for the left, middle and right column, respectively. For the 3 plots in the top row, ammonium sulfate is varied across the y-axis from 0 to 1 M while BME concentration is fixed at 0.5 M and heating time is fixed at 60 minutes. For the 3 graphs in the middle row, BME concentration is varied across the y-axis from 0 to 1 M while ammonium sulfate concentration is fixed at 1 M and heating time is fixed at 60 minutes. For the 3 graphs in the bottom row, heating time is varied across the y-axis from 5 to 60 minutes while ammonium sulfate concentration is fixed at 1 M and BME concentration is fixed at 0.5 M.

Building upon the further improvements using the monomeric benzonase with reducing conditions, we then further optimized precipitation conditions using standard design of experiments (DoE) methodology as illustrated in Figure 3c. The temperature, salt concentration (ammonium sulfate), pH, BME concentration and time for the precipitation were varied and a model built to understand the relationship between these key variables and nuclease removal. A graphical overview of this model is given in Figure 3d. Importantly, numerous combinations of temperature, pH, salt concentration and BME concentrations enable effective removal of nuclease activity. This should allow for extending this methodology to numerous proteins of interest, with varied solubility and isoelectric points and thermal stabilities (at least greater than 60 degrees Celsius).

As Figure 3c-d demonstrates, a range of conditions lead to an optimal precipitation, with the monomeric benzonase being essentially completely removed. We chose the following purification condition: 80°C for 1 hour at pH 9.0 in the presence of 1 M ammonium sulfate and 0.5 M BME; and validated the performance of these Taq preparations in routine PCR compared to commercial Taq. Taq preparations were >95% pure (Figure 4a) and performed equally, if not better, than commercial preparations of Taq (Figure 4b-d). Importantly, these Taq preparations are DNA/RNA free.

**Figure 4.**
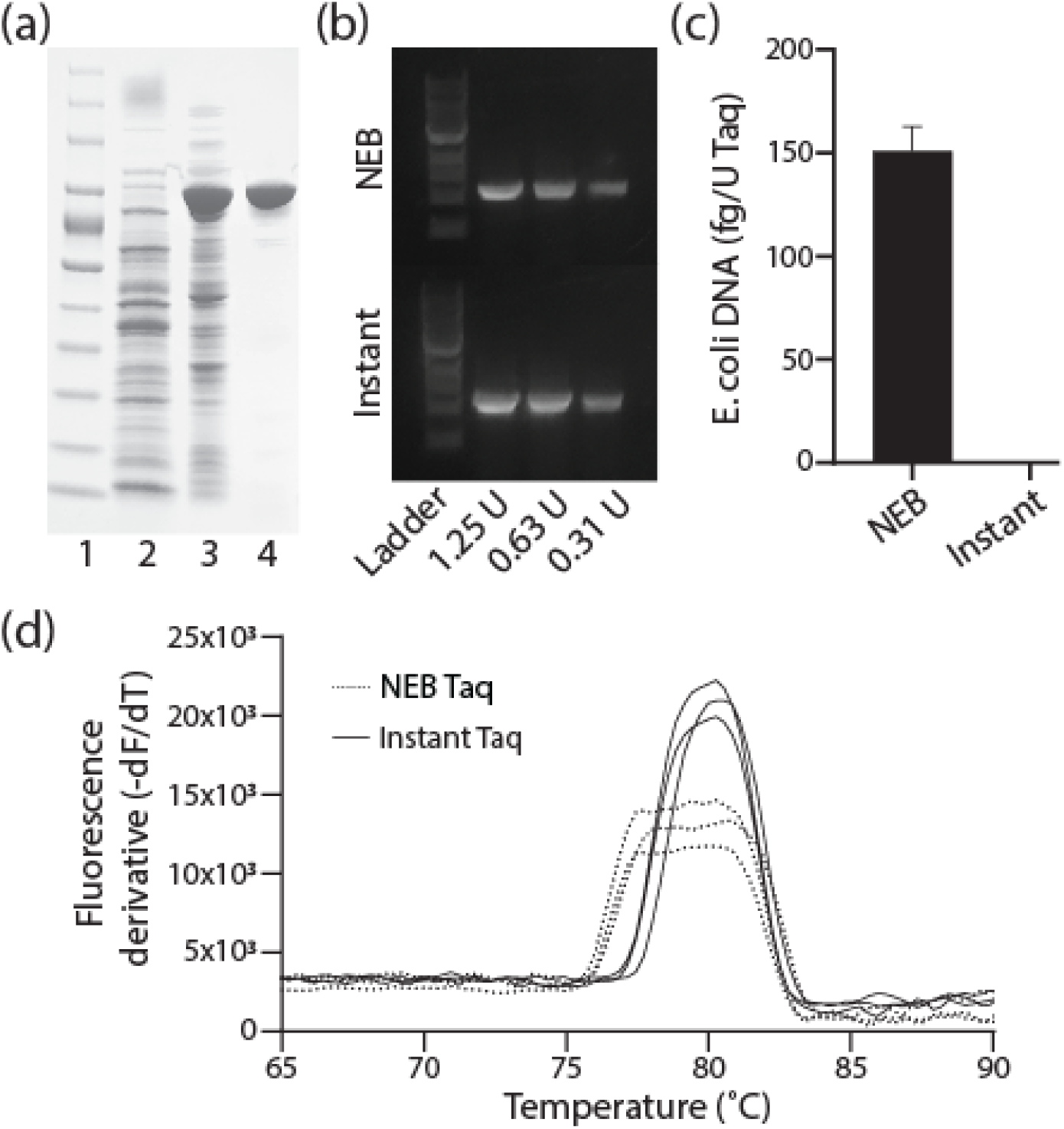
Quality of Instant Taq. (a) Taq Purification using the optimal precipitation condition. Protein ladder (lane 1), Lysate of DLF_R005 with an empty vector control (lane 2), Lysate of DLF_R005 expressing Taq (lane 3) and Taq after optimal precipitation condition (lane 4, 80 °C for 60 minutes at pH 9.0 and 0.5 M beta-mercaptoethanol). (b) Agarose gel image of a PCR product made using NEB Taq (upper panel) or Instant Taq (bottom panel). (c) *E. coli* chromosomal DNA per unit of Taq for NEB Taq and Instant Taq, as measured by qPCR (n=3). (d) qPCR melt curves of a PCR product made using NEB Taq (dotted line) or Instant Taq (solid line), (n=3)

## Discussion

The combination of i) two-stage protein expression, ii) genetic circuits enabling cell autolysis and DNA/RNA autohydrolysis and iii) improved protein precipitation conditions have yielded a rapid, low-cost method for the high titer expression and “Instant” purification of Taq polymerase. The average lab can use this method for making and purifying Taq polymerase in-house with minimal hands-on time or specialized equipment and could lead to significant cost savings. While the purified Taq made using this method is not 99.9+% pure, this is seldom needed for many applications such as routine PCR and cloning, where a relatively pure enzyme (free of key unwanted activities) is sufficient. This method can be adapted to additional DNA modifying enzymes and research proteins.

Additionally, as described previously, two-stage protein expression, upon phosphate depletion, is readily scalable and results in high titers with different proteins with no process optimization ^10^. As these methods are amenable to microtiter plates, the use of this chromatography-free purification approach can further reduce the work required in high throughput screening workflows, such as those used in protein engineering. Generally, this methodology can be leveraged in bioprocess development where product leads can be validated in high throughput screens and easily scaled-up, reducing development timelines.

While very promising, this method is not without its limitations. First, it is currently restricted to thermostable proteins as heat is required to remove nuclease activity. Similarly, using this approach for proteins with oxidized disulfide bonds may not be ideal as BME is required in many cases to completely remove nuclease activity. Conditions for complete nuclease removal at the low temperature end (60 °C) are also more restricted and may not be compatible with some mildly thermostable proteins. Yet, the use of new and more thermolabile nuclease mutants/variants would be expected to increase the range of acceptable conditions.^16^ Exploring additional process variables such as different salts or other additives may very well also expand the approach to a wider array of proteins.

Last, while heat precipitation has been previously extensively used for protein purification, including Taq, additional purification steps are commonly required before the protein of interest can be used. These results highlight that although the ever-expanding arsenal of synthetic biology tools is powerful, the combination of synthetic biology with more traditional approaches to optimization (such as DoE) can be even more so. The combination of new synthetic biology tools with traditional approaches may well yield the most practical solutions.

## Materials & Methods

### Media, Buffers and Reagents

Refer to Table S1 for a complete list of reagents and catalog numbers. LB Lennox formulation was used for routine strain propagation and 5mL starter cultures. Autoinduction Broth (AB) was prepared as described by Menacho-Melgar et al.^10^ Autoinduction Broth 2 (AB-2) was prepared as recently reported.^18^ Buffer compositions are as follows: Lysis Buffer (20 mM Tris, pH 8.0, 0.1% Triton-X 100, supplemented with a protein inhibitor cocktail), 2× Precipitation Buffer (1 M Tris-HCl, pH 9.0, 2 M NH_4_SO_4_), 2× Storage buffer (40 mM of Tris-HCl, pH 8, 100 mM KCl, 0.1% Nonidet P40 Substitute, 0.1% Tween-20, 0.1 mM EDTA, 50% glycerol), NEB 10x Standard Taq Reaction buffer (100 mM Tris-HCl, 500 mM KCl, 15 mM MgCl_2_, pH 8.3) (New England Biolabs, Ipswich, MA), 2× PCR reaction mix (2× Standard Taq Reaction buffer, 0.4 mM dNTPs), 2× Loading Buffer (4% w/v sodium dodecyl sulfate, 4 mM beta-mercaptoethanol, 8% v/v glycerol, 80 mM Tris-HCl, pH 6.8, 0.02% w/v bromophenol blue)

### Plasmids & Strains

Strain DLF_R004 was constructed as previously reported.^11^ Strain DLF_R005 was similarly constructed.^19^ Briefly, a double stranded synthetic DNA, “del_ompT-lysozyme-benz-H184R”, was obtained (gBlock™, IDT, Coralville, IA) and integrated into the chromosome using standard recombineering methods. This gBlock™ codes an operon expressing lambda phage lysozyme and a benzonase™ mutant (H184R) under the control of the low-phosphate inducible promoter yibDp, and contains flanking homology sequences to the *ompT* gene. Chromosomal integration was confirmed using Sanger sequencing (Genewiz, Morrisville, NC). The recombineering plasmid pSIM5 was a kind gift from Donal Court (NCI, https://redrecombineering.ncifcrf.gov/court-lab.html). Synthetic DNA sequences including gBlock™ and oligonucleotides are given in Table S2.

Plasmid pTwistKan-yibDp-Taq is a high copy plasmid, conferring kanamycin resistance and enabling the low phosphate induction of Taq from the robust *yibD* gene promoter.^9^ This plasmid, was constructed as sequence verified clones by Twist Bioscience (San Francisco, CA). Plasmid pHCKan-yibDp-GFP was constructed as described previously.^10^ Plasmid pHCKan-yibDp-tsGFP was assembled from a gBlock™ and store-bought linearized pHCKan plasmid (Lucigen, Middleton, WI) using NEBuilder® HiFi DNA Assembly Mix (New England Biolabs, Ipswich, MA) according to the manufacturer’s instructions. Correct assembly was confirmed using Sanger sequencing (Genewiz, Morrisville, NC). Plasmid sequences are given in Table S3. Plasmid pLJM1-EGFP was a generous gift from David Sabatini (Addgene plasmid #19319).

### Autoinduction (Hands on Time < 10 minutes)

A colony or cryostock of strain DLF_R004 or DLF_R005 containing the plasmid coding Taq is used to start a 5mL culture of LB (Lennox formulation) with 35 μg/mL kanamycin. This starter culture is grown overnight (~16 h) at 37 °C in a shaking incubator. Then, 200 μL of the starter culture is used to inoculate 20 mL of either Autoinduction Broth (AB) or (AB2)^10,14^ (with 35 μg/mL kanamycin) in a 250 mL baffled Erlenmeyer flask. This culture is incubated at 37 °C shaking at 150 rpm for 24 hrs.

### Autolysis & Auto DNA/RNA hydrolysis (Hands on Time < 10 minutes)

After 24 hours in AB, cells are harvested by centrifugation and resuspended in Lysis buffer using 1/10th of the original volume (2 mL for a single 20 mL culture in a 250 mL flask). The lysate can be aliquoted into 1.5 mL microcentrifuge tubes to facilitate lysis. Resuspended pellets are frozen for 60 minutes and then incubated at 60°C for one hour followed by a two hour incubation at 37 °C to complete lysis and auto DNA/RNA hydrolysis. After lysis, samples were centrifuged at 14,000 rcf for one minute in a microcentrifuge.

### Heat Precipitation (Hands on Time < 5 minutes)

The lysate is mixed with 1 volume of 2× Precipitation buffer (1 ml of 2x buffer per 1 ml of lysate). Beta-mercaptoethanol is added to the mixture to a final 0.5 M concentration and heated to 80°C for 60 minutes after which lysates are again clarified by centrifugation at 13,000 rpm (15,000 g) for two minutes in a microcentrifuge. The purified Taq solution is then mixed with 1 volume of 2× Storage buffer and stored at −20 °C.

### Routine PCR

Taq in storage buffer can be used for routine PCR by adding 1 μL of Taq (or 1.25 units) to a 50 μL PCR reaction, along with 25 μL of 2× PCR Reaction Mix, 1 μL of template DNA, 2.5 μL of each primer (5-10 μM stock concentration of primers) and 19 μL of molecular biology grade water. In this study, all PCR products shown in an agarose gel were made using pHCKan-yibDp-GFP as template and SL1 and SR2 oligonucleotides as primers. For these, an annealing temperature of 53 °C and extension time of 2 minutes was used.

### Design of experiments

Shake-flask cell cultures were prepared and grown as described above. To ensure constant protein levels across experiments, cells were lysed in 1/50th of the original culture volume in Lysis buffer. After lysis, protein concentration was measured and diluted to 10 μg/μl. To perform the DoE, 9 different pH and ammonium concentration combinations were required. For each combination, a separate solution containing 1 M buffer was prepared where MES, MOPS and Tris buffers were used in buffers of pH 6, pH 7.5 and pH 9, respectively. For all experiments, beta-mercaptoethanol was added before the precipitation step. All experiments were performed using a thermocycler (Bulldog Bio, Portsmouth, NH). The experimental design was a definitive screening design and was tabulated using JMP Pro 15 software (SAS Institute, Cary, NC). The design was later augmented to better estimate higher order factors while minimizing confounding effects. The model was constructed iteratively, maximizing the parameters: R-squared, adjusted R-squared and Press.

### Nuclease assays

To rapidly determine if there was residual nuclease activity in lysates, 200 μg of the plasmid, pHCKan-yibDp-GFP, was incubated with 1 μl of lysate in a 10 μl reaction volume. After incubation at 37 °C for 30 minutes, the samples were run in an agarose gel. To assess residual nuclease activity in purified TAQ preparations, a routine PCR was performed as described above and run in an agarose gel immediately after the PCR was completed and after incubation at 4 °C overnight. For absolute quantification, nuclease activity was measured using DNAseAlert™ kit (IDT, Coralville IA). Briefly, 2 nmol of substrate was resuspended in 200 μl nuclease-free water. For each reaction, 5 μl of each standard or sample was mixed with 5 μl of resuspended substrate, 5 μl of 10x DNAseAlert™ buffer, and 35 μl nuclease-free water. The mix was placed in a 384-well black plate (Ref# 781090)(Greiner, Monroe, NC) and incubated at 37 °C for 4 hours. Fluorescence was measured using a Tecan Infinite 200 plate reader at 530/560 nm excitation/emission wavelengths. Purchased benzonase (VWR, Radnor, PA) at 78, 39, 19.5, 9.7 and 0 μunits/μl were used as standards.

### qPCR

qPCR melt curves were obtained using an Open qPCR thermocycler (Chai Bio, Santa Clara, CA) and using plasmid pLJM1-EGFP as a template and oligonucleotides pLJM1-4_F and pLJM1-4_R as primers (sequences in Table S2). The PCR mix was prepared as described above with addition of 0.5x of thiazole green. 45 cycles of 10 s at 95 °C followed by 30 s at 60 °C were completed before obtaining the melt curve. For the melt curve, the temperature was ramped from 60 °C to 95 °C at a rate of 0.09 °C/s.

### E. coli chromosomal DNA quantification

Chromosomal DNA was quantified using an Open qPCR thermocycler. *E. coli* DNA standards were purchased (Fisher Scientific, Waltham, MA) and used at 10-fold dilutions between 2000 pg/μl and 0.002 pg/μl. Chrom_F and Chrom_R were used as primers and Chrom_probe as the reaction probe (sequences in Table S2). First, a 10x primer/probe mix was prepared by combining the primers with the probe at a concentration of 5 μM and 2.5 μM, respectively. Each reaction consisted of 10 μl of Luna Universal Probe qPCR Master Mix (New England Biolabs, Ipswich, MA), 5 μl of standard or sample, 2 μl of 10x primer/probe mix and 3 μl of nuclease-free water. The qPCR consisted of 45 cycles of 10 s at 95 °C followed by 30 s at 60 °C.

### Protein Quantification

Protein was quantified using Bradford reagent and albumin standards (Thermo Scientific, Waltham, MA). Bradford reagent was made by dissolving 50 mg of Coomassie Blue G250 (Biobasic, Amherst, NY) in 50 ml of methanol then mixed with 100 ml of 85% phosphoric acid. After adding 850 ml of ultrapure water, the solution was filtered using a coffee filter and stored at 4 °C. To perform the Bradford assay, 5 μl of standard or sample was mixed with 150 μl of Bradford reagent. Absorbance was then measured using a spectrophotometer (Molecular Devices, San Jose, CA) at 595 nm.

### SDS-PAGE

Protein samples were mixed 1:1 with 2× loading buffer. Samples were then heated at 95 °C for 30 s, loaded in a 4-15% gradient Mini-Protean TGX precast protein gel (Bio-Rad Laboratories, Hercules, CA) and ran at 140 V. For lysate samples, 10 μg of total protein were loaded per lane. For purified Taq samples, 3 μg of total protein were loaded per lane.

## Acknowledgements

We would like to acknowledge the following support: ONR YIP #12043956, DOE EERE grant #EE0007563 and North Carolina Biotechnology Center 2018-BIG-6503, NIH R61 AI140485-01.

## Author contributions

R. Menacho-Melgar constructed plasmids and strains, performed expression and purification. T. Yang performed purification studies. M.D. Lynch designed and analyzed experiments. All authors wrote, revised and edited the manuscript.

## Conflicts of Interest

M.D. Lynch has a financial interest in DMC Biotechnologies, Inc. M.D. Lynch, and R. Menacho-Melgar have a financial interest in Roke Biotechnologies, Inc.

