## Supplemental Materials for "Instant Taq: Rapid Autoinducible Expression and Chromatography-free Purification of Taq polymerase"

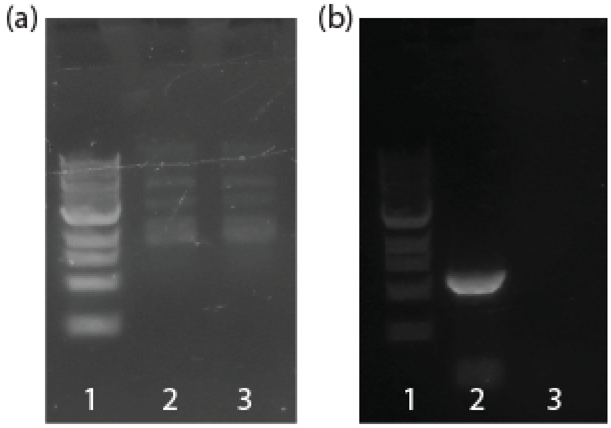

Figure S1. Taq activity loss after treatment with 0.1 M NaOH and 70 °C for 30 minutes. (a) Removal of nuclease activity as shown in an agarose gel: DNA ladder (lane 1), pHCKan-yibDp-GFP incubated without (lane 2) or with treated lysate (lane 3). (b) Taq activity loss as shown in an agarose gel: DNA ladder (lane 1), PCR product using commercial Taq (NEB, lane 2) and PCR product using purified Taq in the presence of 0.1 M NaOH (lane 3)

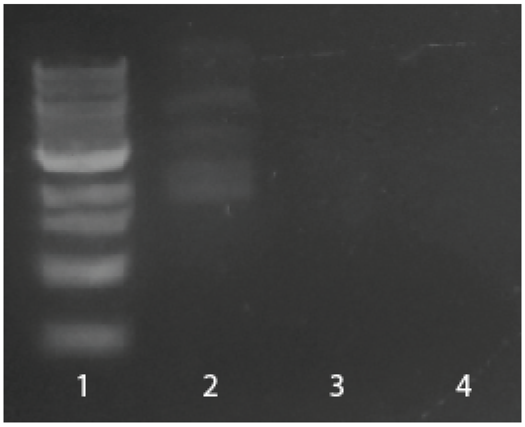

Figure S2. Residual nuclease activity as shown in an agarose gel: DNA ladder (lane 1), pHCKan-yibDp-GFP incubated without lysate (lane 2), and with the upper (lane 3) or lower (lane 4) phase of lysate after aqueous two-phase extraction

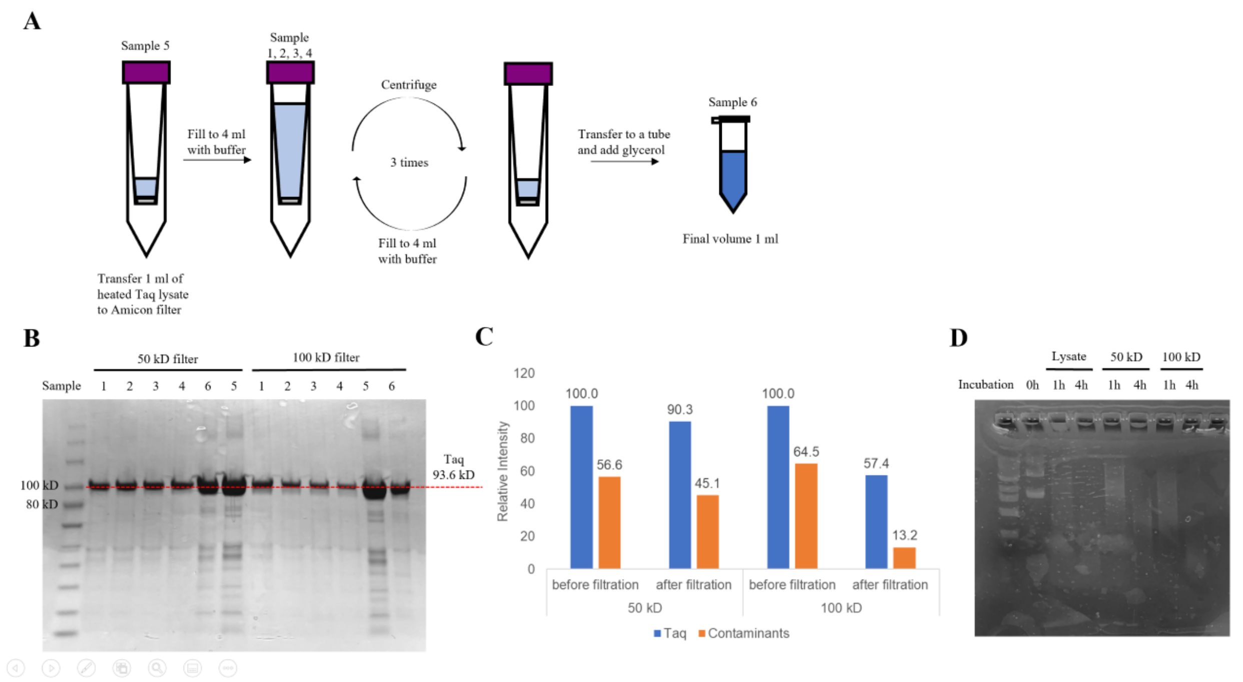

Figure S3. Nuclease activity remained after diafiltration. a)the diafiltration is done via an iterative process of centrifugation using a filter column (with MWCO of 50kD or 100kD) followed by refilling the column with the buffer. b)in each iteration, a sample is taken from the column for SDS-PAGE analysis (Sample 1-4: sample after 0-3 times of refilling buffer, respectively; Sample 5: original lysate; Sample 6: final concentrate). c)densitometry analysis evaluates loss of taq as well as impurity protein after diafiltration. d)original lysate as well as the diafiltrates are subject to plasmid spike-in assay to assess nuclease activity.

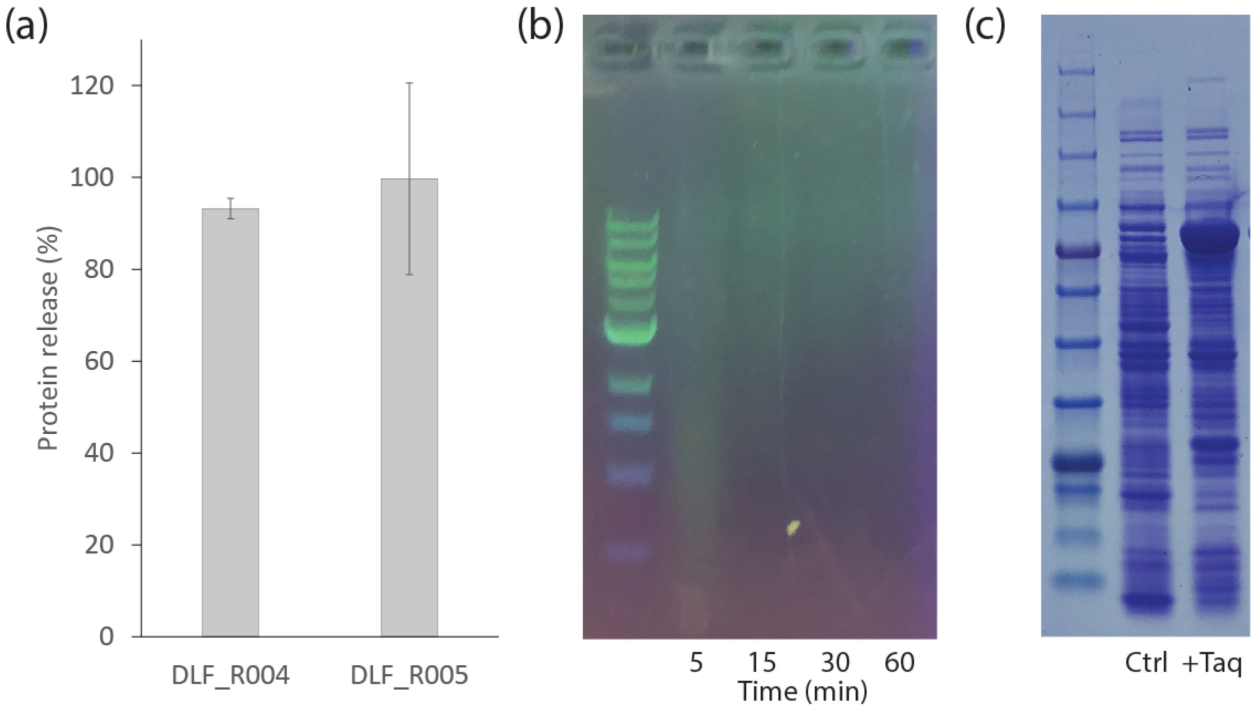

**Figure S4:** DLF_R005 autolysis/autohydrolysis validation. (a) GFP protein release after autolysis using DLF_R005 compared to DLF_R004 (data adapted from (Menacho-Melgar et al. 2020)). (b) Time-course showing DNA/RNA autohydrolysis of DLF_R005 after thawing and incubating at 37 °C for 1 hour. (c) SDS-PAGE gel showing expression of Taq in DLF_R005 (lane 3) compared to DLF_R005 with empty vector control (lane 2).

### Table S1: Reagents used in this study

| **Reagent** | **Supplier** | **Catalog Number** |
|---|---|---|
| Tryptone | Biobasic | TG217(G211) |
| Yeast extract | Biobasic | G0961 |
| Sodium chloride | Biobasic | DB0483 |
| Ammonium sulfate anhydrous | VWR | M105 |
| Citric acid | Chem-Impex International | 02107 |
| 18 M sulfuric acid | VWR | BDH3096 |
| Cobalt sulfate heptahydrate | Acros Organics | 213101000 |
| Copper sulfate pentahydrate | Alfa-aesar | 14178 |
| Zinc sulfate heptahydrate | Sigma Aldrich | 307491 |

| | | |
|---|---|---|
| Sodium molybdate dihydrate | Alfa-aesar | 12214 |
| Boric acid | Spectrum chemical | B1125 |
| Manganese sulfate monohydrate | Alfa-aesar | 33341 |
| Iron (II) sulfate heptahydrate | Sigma Aldrich | 215422 |
| Magnesium sulfate heptahydrate | Caisson labs | M020 |
| Calcium sulfate dihydrate | Alfa-aesar | 36700 |
| Potassium sulfate monobasic | Biobasic | PRB0445 |
| Potassium sulfate dibasic | Biobasic | PRB0447 |
| Glucose | VWR | 89405-376 |
| 3-morpholinopropane-1-sulfonic acid (MOPS) | GoldBio | M-790-1 |
| Bis-Tris | GoldBio | B-020-500 |
| Thiamine hydrochloride | MP Biomedicals | 103028 |
| Casamino acids | Biobasic | CB3060 |
| 12 M hydrochloric acid | Biobasic | HC6025 |
| Tris base | GoldBio | T-400-500 |
| Triton X-100 | Sigma Aldrich | 93443 |
| Protease inhibitors tablets EDTA-free | Thermofisher Scientific | A32965 |
| Nonidet P40 substitute | Sigma Aldrich | 74385 |
| Tween-20 | Sigma Aldrich | P9416 |
| EDTA | Fisher Chemical | 5311-100 |
| Glycerol | Reagents | C1210500-4C |
| Potassium chloride | Alfa-aesar | J64189 |
| Magnesium chloride | Biobasic | MRB0328 |
| dNTPs | New England Biolabs | N0447S |
| Beta-mercaptoethanol | Sigma Aldrich | M6250-1L |
| 2-(N-morpholino)ethanesulfonic acid (MES) | Biobasic | MB0341 |
| Taq polymerase | New England Biolabs | M0273S |

| | | |
|---|---|---|
| 10x Taq reaction buffer | New England Biolabs | B9014S |
| Benzonase | VWR | 10011-072 |
| 2x Luna Universal qPCR reaction mix | New England Biolabs | M3003S |
| 10,000x Thiazole green | Biotum | 40086-0.5 |
| Albumin standards | ThermoFisher Scientific | 23209 |
| Coomassie blue G-250 | Biobasic | CB0038 |
| Phosphoric acid | VWR | UN180J |
| Sodium dodecyl sulfate | Fisher Scientific | BP166-500 |
| Bromophenol blue | Biobasic | BB2230 |

### Table S2. Synthetic DNA and oligonucleotides used in this study

| **del_ompT-lysozyme-benz-H184R** |
|---|
| ATGGATAACAAAAATATCACTCAGGTGGCGCAATTATGTGGCTATAGCAGCACGTCGTACTTTATCTCT<br>GTTTTTAAGGCGTTTTACGGCCTGACACCGTTGAATTATCTCGCCAAACAGCGACAAAAAGTGATGTG<br>GtgaAGGGCAAAGCGGAAACGGATAAGACGGGCATAAATGAGGAAGAAATGGCTCGACCTAGCATAA<br>CCCCGCGGGGCCTCTTCGGGGGTCTCGCGGGGTTTTTTGCTGAAAGAAGCTTCAAATAAAACGAAAG<br>GCTCAGTCGAAACACTGGGCCTTTCGTTTTATCTGTTGTTTGTCGCTGCGGCCGGGTCAGGTATGATTT<br>AAATGGTCAGTAACGGGTCTTGAGGGGTTTTTTGCATATGTGCGTAATTGTGCTGATCTCTTATATAGC<br>TGCTCTCATTATCTCTCTACCCTGAAGTGACTCTCTCACCTGTAAAAATAATATCTCACAGGCTTAATAG<br>TTTCTTAATACAAAGCCTGTAAAACGTCAGGATAACTTCTGTGTAGGAGGATAATCTTTTGCTGGAAA<br>AAGGAGATATACCATGGTGGAGATCAATAACCAGCGTAAAGCATTTTTGGATATGTTAGCGTGGTCTG<br>AAGGGACTGACAATGGCCGTCAGAAAACACGTAACCATGGCTATGACGTGATCGTGGGCGGTGAATT<br>ATTCACAGACTATTCGGACCATCCACGCAAATTAGTAACCCTTAATCCGAAGCTGAAGTCGACGGGCG<br>CGGGCCGCTATCAACTGCTTAGCCGTTGGTGGGATGCTTACCGCAAGCAGTTAGGGCTGAAGGATTTT<br>TCGCCGAAGAGCCAGGATGCTGTCGCTCTGCAACAAATTAAAGAGCGTGGGGCTTTACCAATGATCG<br>ACCGTGGTGACATCCGTCAGGCCATTGATCGCTGCTCTAACATCTGGGCATCTCTTCCAGGTGCGGGC<br>TACGGGCAGTTCGAGCACAAAGCCGACTCCCTTATTGCTAAGTTCAAAGAAGCGGGAGGCACGGTTC<br>GCGAGATCGATGTATAAGGATCTAGGAGGGAGATCATATGCGTTTTAATAATAAAATGTTAGCTCTTGC<br>CGCATTATTGTTTGCAGCCCAAGCGTCAGCGGATACGCTTGAATCTATCGATAACTGTGCCGTGGGTTG<br>CCCGACCGGGGGTTCGTCTAATGTAAGCATCGTCCGTCACGCCTATACCCTTAACAATAATTCTACCAC<br>GAAATTCGCGAACTGGGTGGCCTACCATATTACCAAAGATACTCCCGCATCGGGTAAAACACGCAACT<br>GGAAGACTGACCCGGCACTGAATCCAGCCGATACATTGGCTCCGGCAGATTACACAGGTGCAAACGC<br>TGCTCTGAAAGTAGACCGCGGACATCAGGCTCCGCTTGCGAGTCTGGCTGGAGTTAGTGACTGGGAA<br>TCACTGAACTATTTGTCGAATATTACCCCACAGAAATCTGACTTGAATCAAGGCGCGTGGGCTCGTTT<br>AGAAGACCAAGAGCGTAAACTTATCGACCGTGCAGACATTTCCAGCGTCTATACAGTAACAGGTCCT<br>CTGTACGAGCGCGACATGGGAAAGCTTCCTGGTACTCAGAAAGCTCATACAATCCCTTCTGCATATTG<br>GAAGGTTATCTTTATTAATAATAGCCCGGCTGTCAATCGTTATGCCGCCTTCTTATTTGATCAGAACACT<br>CCCAAGGGGGCTGACTTCTGCCAATTCCGCGTCACTGTAGACGAGATTGAGAAGCGCACGGGGTTAA<br>TCATCTGGGCCGGATTACCTGACGATGTGCAGGCTTCCTTGAAATCCAAGCCGGGAGTTCTTCCCGAG<br>TTGATGGGTTGCAAAAACTAATAAACCGTGTTGACAATTAATCATCGGCATAGTATATCGGCATAGTAT<br>AATACGACAAGGTGAGGAACTAAACCatgtcatcagcggtggagtgcaatgtcgtgcaatacgaatggcgaaaagccgagctcatcggtca<br>gcttctcaaccttggggttacccccggcggtgtgctgctggtccacagctccttccgtagcgtccggcccctcgaagatgggccacttggactgatcgaggccctgc<br>gtgctgcgctgggtccgggagggacgctcgtcatgccctcgtggtcaggtctggacgacgagccgttcgatcctgccacgtcgcccgttacaccggaccttggagtt |

| |
|---|
| gtctctgacacattctggcgcctgccaaatgtaaagcgcagcgcccatccatttgcctttgcggcagcggggccacaggcagagcagatcatctctgatccattgccc<br>ctgccacctcactcgcctgcaagcccggtcgcccgtgtccatgaactcgatgggcaggtacttctcctcggcgtgggacacgatgccaacacgacgctgcatcttgc<br>cgagttgatggcaaaggttccctatggggtgccgagacactgcaccattcttcaggatggcaagttggtacgcgtcgattatctcgagaatgaccactgctgtgagcg<br>ctttgccttggcggacaggtggctcaaggagaagagccttcagaaggaaggtccagtcggtcatgcctttgctcggttgatccgctcccgcgacattgtggcgacag<br>ccctgggtcaactgggccgagatccgttgatcttcctgcatccgccagaggcgggatgcgaagaatgcgatgccgctcgccagtcgattggctgagctcaGAAC<br>GCCAACTAAAATTTCCCCGAGGTGAAAATCGCCCCGGGGAATAACTAGCCATTTCAATGTAACAATTA<br>ACCCTTAAAATAAACCCAGAAGGTTATTAACTAAATCACATAGAAAACCATCAATTATAGTATGTATAA<br>AATAGGCGACAGCAACCCAATTAC |
| **OmpT_dn** |
| GATTATTATGGTGTCACGCCATCTC |
| **OmpT_1000bpup** |
| GCATTGCTTTTTACCGTATTGTCTAAC |
| **benzostrain_seq_conf_2** |
| ATGGTGGAGATCAATAACCAGCGTAAAGC |
| **benzostrain_seq_conf_3** |
| TAGCTCTTGCCGCATTATTGTTTGC |
| **benzostrain_seq_conf_4** |
| CCTTCTTATTTGATCAGAACACTC |
| **SL1** |
| CAGTCCAGTTACGCTGGAGTC |
| **SR2** |
| GGTCAGGTATGATTTAAATGGTCAGT |
| **pLJM1-4_F** |
| GATAGTAGGAGGCTTGGTAGGT |
| **pLJM1-4_R** |
| AGGTGGGTCTGAAACGATAATG |
| **Chrom_F** |
| CCTTACGACCAGGGCTACACA |
| **Chrom_R** |
| CTCGCGAGGTCGCTTCTC |
| **Chrom_Probe** |

| |
|---|
| 5′-FAM520-CGTGCTACAATGGCGCATACA-ZEN/IBLK-3′ |

**Table S3: Plasmid Sequences**

| **pHCKan-yibDp-Taq** |
|---|
| AGGCTAGGTGGAGGCTCAGTGATGATAAGTCTGCGATGGTGGATGCATGTGTCATGGTCATAGCTGTT<br>TCCTGTGTGAAATTGTTATCCGCTCAGAGGGCACAATCCTATTCCGCGCTATCCGACAATCTCCAAGAC<br>ATTAGGTGGAGTTCAGTTCGGCGTATGGCATATGTCGCTGGAAAGAACATGTGAGCAAAAGGCCAGC<br>AAAAGGCCAGGAACCGTAAAAAGGCCGCGTTGCTGGCGTTTTTCCATAGGCTCCGCCCCCCTGACGA<br>GCATCACAAAAATCGACGCTCAAGTCAGAGGTGGCGAAACCCGACAGGACTATAAAGATACCAGGC<br>GTTTCCCCCTGGAAGCTCCCTCGTGCGCTCTCCTGTTCCGACCCTGCCGCTTACCGGATACCTGTCCG<br>CCTTTCTCCCTTCGGGAAGCGTGGCGCTTTCTCATAGCTCACGCTGTAGGTATCTCAGTTCGGTGTAG<br>GTCGTTCGCTCCAAGCTGGGCTGTGTGCACGAACCCCCCGTTCAGCCCGACCGCTGCGCCTTATCCG<br>GTAACTATCGTCTTGAGTCCAACCCGGTAAGACACGACTTATCGCCACTGGCAGCAGCCACTGGTAAC<br>AGGATTAGCAGAGCGAGGTATGTAGGCGGTGCTACAGAGTTCTTGAAGTGGTGGCCTAACTACGGCT<br>ACACTAGAAGAACAGTATTTGGTATCTGCGCTCTGCTGAAGCCAGTTACCTTCGGAAAAAGAGTTGG<br>TAGCTCTTGATCCGGCAAACAAACCACCGCTGGTAGCGGTGGTTTTTTTGTTTGCAAGCAGCAGATTA<br>CGCGCAGAAAAAAAGGATCTCAAGAAGATCCTTTGATCTTTTCTACGGGGTCTGACGCTCTATTCAAC<br>AAAGCCGCCGTCCCGTCAAGTCAGCGTAAATGGGTAGGGGGCTTCAAATCGTCCGCTCTGCCAGTGT<br>TACAACCAATTAACAAATTCTGATTAGAAAAACTCATCGAGCATCAAATGAAACTGCAATTTATTCATA<br>TCAGGATTATCAATACCATATTTTTGAAAAAGCCGTTTCTGTAATGAAGGAGAAAACTCACCGAGGCA<br>GTTCCATAGGATGGCAAGATCCTGGTATCGGTCTGCGATTCCGACTCGTCCAACATCAATACAACCTAT<br>TAATTTCCCCTCGTCAAAAATAAGGTTATCAAGTGAGAAATCACCATGAGTGACGACTGAATCCGGTG<br>AGAATGGCAAAAGCTTATGCATTTCTTTCCAGACTTGTTCAACAGGCCAGCCATTACGCTCGTCATCA<br>AAATCACTCGCATCAACCAAACCGTTATTCATTCGTGATTGCGCCTGAGCGAGACGAAATACGCGATC<br>GCTGTTAAAAGGACAATTACAAACAGGAATCGAATGCAACCGGCGCAGGAACACTGCCAGCGCATC<br>AACAATATTTTCACCTGAATCAGGATATTCTTCTAATACCTGGAATGCTGTTTTCCCGGGGATCGCAGT<br>GGTGAGTAACCATGCATCATCAGGAGTACGGATAAAATGCTTGATGGTCGGAAGAGGCATAAATTCCG<br>TCAGCCAGTTTAGTCTGACCATCTCATCTGTAACATCATTGGCAACGCTACCTTTGCCATGTTTCAGAA<br>ACAACTCTGGCGCATCGGGCTTCCCATACAATCGATAGATTGTCGCACCTGATTGCCCGACATTATCGC<br>GAGCCCATTTATACCCATATAAATCAGCATCCATGTTGGAATTTAATCGCGGCCTCGAGCAAGACGTTT<br>CCCGTTGAATATGGCTCATAACACCCCTTGTATTACTGTTTATGTAAGCAGACAGTTTTATTGTTCATGA<br>TGATATATTTTTATCTTGTGCAATGTAACATCAGAGATTTTGAGACACAACGTGGCTTTCCCCCGCCGC<br>TCTAGAACTAGTGGATCCAAATAAAACGAAAGGCTCAGTCGAAAGACTGGGCCTTTCGTTTTATCTGT<br>TGTTTGTCGCATTATACGAGACGTCCAGGTTGGGATACCTGAAACAAAACCCATCGTACGGCCAAGG<br>AAGTCTCCAATAACTGTGATCCACCACAAGCGCCAGGGTTTTCCCAGTCACGACGTTGTAAAACGAC<br>GGCCAGTCATGCATAATCCGCACGCATCTGGAATAAGGAAGTGCCATTCCGCCTGACCTTGCCCAGGC<br>ATCAAATAAAACGAAAGGCTCAGTCGAAAGACTGGGCCTTTCGTTTTATCTGTTGTTTGTCGGTGAAC<br>GCTCTCTACTAGAGTCACACTGGCTCACCTTCGGGTGGGCCTTTCTGCGTTTATACACAGCTAACACC<br>ACGTCGTCCCTATCTGCTGCCCTAGGTCTATGAGTGGTTGCTGGATAACGTGCGTAATTGTGCTGATCT<br>CTTATATAGCTGCTCTCATTATCTCTCTACCCTGAAGTGACTCTCTCACCTGTAAAAATAATATCTCACA |

GGCTTAATAGTTTCTTAATACAAAGCCTGTAAAACGTCAGGATAACTTCTATATTCAGGGAGACCACA
ACGGTTTCCCTCTACAAATAATTTTGTTTAACTTTCGTGTGTAGGAGGATAATCTATGCTTCCTTTATTC
GAGCCTAAAGGCCGTGTACTGTTAGTTGACGGGCACCACCTTGCTTACCGTACGTTCCACGCCCTTAA
AGGGCTGACAACATCGCGTGGAGAACCAGTGCAGGCGGTTTATGGGTTCGCTAAAAGCTTACTGAAA
GCGTTGAAGGAAGATGGGGATGCCGTCATCGTCGTGTTCGACGCGAAAGCTCCATCTTTCCGCCACG
AAGCTTACGGTGGGTACAAGGCAGGTCGTGCGCCTACCCCTGAGGACTTTCCACGTCAGTTAGCATT
AATTAAGGAGTTGGTAGATTTGTTGGGTCTGGCCCGCCTGGAGGTGCCGGGATACGAGGCCGATGAT
GTGCTTGCAAGCCTTGCGAAAAAAGCCGAAAAGGAAGGTTACGAAGTGCGTATTCTTACCGCTGATA
AGGACCTTTATCAGTTGCTGAGTGACCGCATTCATGTTCTTCACCCCGAAGGTTATCTGATCACACCA
GCATGGTTATGGGAAAAGTATGGACTGCGCCCCGATCAATGGGCAGACTATCGTGCTCTTACGGGCGA
TGAATCGGATAACCTGCCTGGAGTCAAAGGGATTGGTGAGAAAACTGCACGTAAATTGTTAGAGGAG
TGGGGCAGTTTGGAGGCGCTGCTTAAAAACTTAGATCGTCTTAAACCTGCCATTCGCGAGAAAATTTT
AGCTCACATGGACGATCTTAAGCTTTCCTGGGACTTAGCTAAGGTTCGCACGGATTTACCACTGGAGG
TCGATTTCGCCAAGCGTCGCGAGCCGGATCGCGAGCGTCTTCGCGCCTTCCTGGAGCGCTTAGAGTTT
GGTTCATTGCTTCACGAGTTTGGGCTTTTAGAGTCGCCGAAGGCTTTAGAGGAAGCTCCTTGGCCGCC
GCCGGAGGGCGCGTTCGTTGGCTTCGTTCTTTCGCGTAAAGAGCCGATGTGGGCTGATTTGTTAGCAT
TAGCGGCAGCTCGTGGAGGGCGTGTTCATCGCGCGCCAGAACCATACAAGGCATTGCGCGACTTAAA
AGAAGCTCGTGGACTTCTGGCCAAGGATTTGTCAGTGCTTGCCTTGCGTGAAGGGCTTGGACTTCCG
CCCGGCGACGACCCGATGTTACTTGCTTATCTGCTGGATCCGTCGAATACAACGCCCGAGGGGGTTGC
GCGTCGTTACGGCGGCGAATGGACTGAGGAAGCGGGGGAACGTGCGGCGCTGTCGGAGCGTTTATTT
GCTAATTTATGGGGTCGTTTGGAGGGTGAAGAACGTCTGCTTTGGTTATATCGCGAAGTCGAGCGTCC
TCTGTCCGCGGTATTAGCTCATATGGAGGCTACAGGGGTCCGTCTTGATGTCGCGTACCTTCGTGCATT
ATCTCTGGAGGTAGCGGAAGAAATCGCCCGTTTGGAAGCCGAAGTGTTTCGCCTGGCCGGACACCCG
TTTAATTTGAACTCGCGTGACCAACTGGAGCGTGTTTTGTTCGACGAACTGGGGTTACCGGCCATCGG
GAAAACTGAGAAGACAGGAAAGCGCAGCACTTCAGCGGCAGTGTTGGAGGCACTGCGTGAGGCGC
ACCCCATCGTTGAAAAGATCTTACAGTACCGTGAACTGACGAAGTTAAAGAGTACCTATATTGACCCC
CTGCCAGATCTTATCCACCCACGTACAGGCCGTTTGCACACGCGCTTCAATCAAACCGCTACTGCAAC
AGGACGTCTTTCTTCTTCTGATCCTAATCTGCAAAATATCCCCGTGCGCACGCCTCTTGGCCAACGCAT
CCGTCGTGCATTCATTGCAGAAGAGGGTTGGTTGCTTGTGGCACTGGACTATTCACAAATCGAGCTTC
GTGTCCTTGCACATTTGTCAGGAGACGAGAACCTGATTCGTGTGTTCCAGGAGGGACGCGACATTCA
CACCGAGACAGCGTCATGGATGTTTGGTGTGCCCCGCGAGGCTGTCGATCCATTAATGCGTCGTGCAG
CGAAGACGATCAACTTTGGTGTGCTGTATGGAATGTCCGCTCACCGCCTTTCCCAAGAGTTGGCTATT
CCATATGAAGAAGCGCAGGCGTTTATTGAACGCTACTTTCAAAGTTTTCCTAAGGTTCGTGCATGGAT
TGAGAAGACTTTAGAAGAGGGGCGCCGTCGCGGTTACGTCGAAACACTGTTTGGCCGCCGCCGTTAT
GTTCCTGACTTAGAGGCACGTGTTAAGTCAGTGCGCGAAGCAGCCGAGCGTATGGCCTTTAACATGC
CCGTACAGGGTACTGCTGCCGACTTGATGAAATTGGCAATGGTCAAGCTGTTTCCGCGTCTGGAAGA
GATGGGGGCCCGTATGTTATTGCAAGTCCACGACGAGTTGGTTCTGGAAGCGCCCAAAGAGCGCGCT
GAGGCGGTGGCGCGTTTGGCGAAGGAGGTGATGGAAGGAGTTTATCCGCTGGCAGTGCCGTTGGAG
GTCGAGGTGGGAATCGGCGAAGACTGGTTATCAGCAAAAGAATAACCGGCTTATCGGTCAGTTTCAC
CTGATTTACGTAAAAACCCGCTTCGGCGGGTTTTTGCTTTTGGAGGGGCAGAAAGATGAATGACTGTC
CACGACGCTATACCCAAAAGAAA

**pHCKan-yibDp-GFP**

TGCCCAGGCATCAAATAAAACGAAAGGCTCAGTCGAAAGACTGGGCCTTTCGTTTTATCTGTTGTTTG
TCGGTGAACGCTCTCTACTAGAGTCACACTGGCTCACCTTCGGGTGGGCCTTTCTGCGTTTATACACA
GCTAACACCACGTCGTCCCTATCTGCTGCCCTAGGTCTATGAGTGGTTGCTGGATAACGTGCGTAATTG

TGCTGATCTCTTATATAGCTGCTCTCATTATCTCTCTACCCTGAAGTGACTCTCTCACCTGTAAAAATAA
TATCTCACAGGCTTAATAGTTTCTTAATACAAAGCCTGTAAAACGTCAGGATAACTTCTATATTCAGGG
AGACCACAACGGTTTCCCTCTACAAATAATTTTGTTTAACTTTCGTGTGTAGGAGGATAATCTATGGCT
AGCAAAGGAGAAGAACTTTTCACTGGAGTTGTCCCAATTCTTGTTGAATTAGATGGTGATGTTAATGG
GCACAAATTTTCTGTCAGTGGAGAGGGTGAAGGTGATGCTACATACGGAAAGCTTACCCTTAAATTTA
TTTGCACTACTGGAAAACTACCTGTTCCATGGCCAACACTTGTCACTACTTTCTCTTATGGTGTTCAAT
GCTTTTCCCGTTATCCGGATCATATGAAACGGCATGACTTTTTCAAGAGTGCCATGCCCGAAGGTTATG
TACAGGAACGCACTATATCTTTCAAAGATGACGGGAACTACAAGACGCGTGCTGAAGTCAAGTTTGA
AGGTGATACCCTTGTTAATCGTATCGAGTTAAAAGGTATTGATTTTAAAGAAGATGGAAACATTCTCGG
ACACAAACTCGAGTACAACTATAACTCACACAATGTATACATCACGGCAGACAAACAAAAGAATGGA
ATCAAAGCTAACTTCAAAATTCGCCACAACATTGAAGATGGATCCGTTCAACTAGCAGACCATTATCA
ACAAAATACTCCAATTGGCGATGGCCCTGTCCTTTTACCAGACAACCATTACCTGTCGACACAATCTG
CCCTTTCGAAAGATCCCAACGAAAAGCGTGACCACATGGTCCTTCTTGAGTTTGTAACTGCTGCTGG
GATTACACATGGCATGGATGAGCTCTACAAATAATGAGGATCCCCGGCTTATCGGTCAGTTTCACCTGA
TTTACGTAAAAACCCGCTTCGGCGGGTTTTTGCTTTTGGAGGGGCAGAAAGATGAATGACTGTCCAC
GACGCTATACCCAAAAGAAAGACGAATTCTCTAGATATCGCTCAATACTGACCATTTAAATCATACCTG
ACCTCCATAGCAGAAAGTCAAAAGCCTCCGACCGGAGGCTTTTGACTTGATCGGCACGTAAGAGGTT
CCAACTTTCACCATAATGAAATAAGATCACTACCGGGCGTATTTTTTGAGTTATCGAGATTTTCAGGAG
CTAAGGAAGCTAAAATGAGCCATATTCAACGGGAAACGTCTTGCTCGAGGCCGCGATTAAATTCCAA
CATGGATGCTGATTTATATGGGTATAAATGGGCTCGCGATAATGTCGGGCAATCAGGTGCGACAATCTA
TCGATTGTATGGGAAGCCCGATGCGCCAGAGTTGTTTCTGAAACATGGCAAAGGTAGCGTTGCCAAT
GATGTTACAGATGAGATGGTCAGGCTAAACTGGCTGACGGAATTTATGCCTCTTCCGACCATCAAGCA
TTTTATCCGTACTCCTGATGATGCATGGTTACTCACCACTGCGATCCCAGGGAAAACAGCATTCCAGGT
ATTAGAAGAATATCCTGATTCAGGTGAAAATATTGTTGATGCGCTGGCAGTGTTCCTGCGCCGGTTGC
ATTCGATTCCTGTTTGTAATTGTCCTTTTAACGGCGATCGCGTATTTCGTCTCGCTCAGGCGCAATCAC
GAATGAATAACGGTTTGGTTGGTGCGAGTGATTTTGATGACGAGCGTAATGGCTGGCCTGTTGAACAA
GTCTGGAAAGAAATGCATAAGCTTTTGCCATTCTCACCGGATTCAGTCGTCACTCATGGTGATTTCTCA
CTTGATAACCTTATTTTTGACGAGGGGAAATTAATAGGTTGTATTGATGTTGGACGAGTCGGAATCGCA
GACCGATACCAGGATCTTGCCATCCTATGGAACTGCCTCGGTGAGTTTTCTCCTTCATTACAGAAACG
GCTTTTTCAAAAATATGGTATTGATAATCCTGATATGAATAAATTGCAGTTTCACTTGATGCTCGATGAG
TTTTTCTAATGAGGGCCCAAATGTAATCACCTGGCTCACCTTCGGGTGGGCCTTTCTGCGTTGCTGGC
GTTTTTCCATAGGCTCCGCCCCCCTGACGAGCATCACAAAAATCGATGCTCAAGTCAGAGGTGGCGA
AACCCGACAGGACTATAAAGATACCAGGCGTTTCCCCCTGGAAGCTCCCTCGTGCGCTCTCCTGTTCC
GACCCTGCCGCTTACCGGATACCTGTCCGCCTTTCTCCCTTCGGGAAGCGTGGCGCTTTCTCATAGCT
CACGCTGTAGGTATCTCAGTTCGGTGTAGGTCGTTCGCTCCAAGCTGGGCTGTGTGCACGAACCCCCC
GTTCAGCCCGACCGCTGCGCCTTATCCGGTAACTATCGTCTTGAGTCCAACCCGGTAAGACACGACTT
ATCGCCACTGGCAGCAGCCACTGGTAACAGGATTAGCAGAGCGAGGTATGTAGGCGGTGCTACAGAG
TTCTTGAAGTGGTGGCCTAACTACGGCTACACTAGAAGAACAGTATTTGGTATCTGCGCTCTGCTGAA
GCCAGTTACCTCGGAAAAAGAGTTGGTAGCTCTTGATCCGGCAAACAAACCACCGCTGGTAGCGGTG
GTTTTTTTGTTTGCAAGCAGCAGATTACGCGCAGAAAAAAAGGATCTCAAGAAGATCCTTTGATTTTC
TACCGAAGAAAGGCCCACCCGTGAAGGTGAGCCAGTGAGTTGATTGCAGTCCAGTTACGCTGGAGT
CTGAGGCTCGTCCTGAATGATATCAAGCTTGAATTCGTT

| **Table S4.** Design of experiments results | | | | | | |
|---|---|---|---|---|---|---|
| **Condition** | **BME (M)** | **Temp (°C)** | **Time (min)** | **pH** | **NH4SO4 (M)** | **Nuclease (µ units)** |
| 1 | 0.5 | 80 | 60 | 9 | 1 | 0 |
| 2 | 1 | 80 | 60 | 6 | 1 | 0 |
| 3 | 0 | 80 | 60 | 9 | 1 | 0 |
| 4 | 1 | 60 | 60 | 9 | 1 | 0 |
| 5 | 1 | 60 | 32.5 | 9 | 1 | 34.5101 |
| 6 | 1 | 80 | 5 | 6 | 1 | 41 |
| 7 | 0 | 80 | 5 | 9 | 1 | 83.8957 |
| 8 | 1 | 60 | 5 | 7.5 | 1 | 90.2317 |
| 9 | 0.5 | 60 | 5 | 6 | 1 | 103.9245 |
| 10 (n=3) | 0.5 | 70 | 32.5 | 7.5 | 0.5 | 110.812 (+/-13.8) |
| 11 | 1 | 70 | 5 | 9 | 1 | 135.3229 |
| 12 | 1 | 80 | 32.5 | 7.5 | 1 | 187.4893 |
| 13 | 0 | 80 | 32.5 | 6 | 0 | 194.9 |
| 14 | 1 | 60 | 60 | 6 | 1 | 202.4141 |
| 15 | 1 | 80 | 5 | 6 | 0.5 | 237.05 |
| 16 | 0 | 80 | 5 | 6 | 1 | 238 |
| 17 | 1 | 60 | 60 | 6 | 0 | 241.2397 |
| 18 | 0 | 80 | 60 | 6 | 1 | 276.7213 |
| 19 | 0 | 60 | 60 | 6 | 0 | 309.8 |
| 20 | 0.5 | 60 | 5 | 6 | 0 | 311 |
| 21 | 0 | 60 | 60 | 9 | 0.5 | 318.95 |
| 22 | 0.5 | 60 | 60 | 9 | 0 | 324.1005 |
| 23 | 0.5 | 80 | 60 | 6 | 0 | 370.3533 |
| 24 | 1 | 80 | 60 | 9 | 0.5 | 417.2397 |
| 25 | 0 | 60 | 60 | 9 | 1 | 422.75 |

| | | | | | | |
|---|---|---|---|---|---|---|
| 26 | 0 | 80 | 32.5 | 9 | 0 | 470.9 |
| 27 | 0 | 80 | 60 | 7.5 | 0 | 521.15 |
| 28 | 0 | 60 | 60 | 6 | 1 | 625.55 |
| 29 | 1 | 70 | 5 | 6 | 0 | 625.7 |
| 30 | 0 | 80 | 5 | 7.5 | 0 | 720.5 |
| 31 | 0 | 70 | 5 | 6 | 1 | 927.8 |
| 32 | 1 | 60 | 5 | 9 | 0 | 1031 |
| 33 | 1 | 80 | 60 | 6 | 0 | 1158 |
| 34 | 0 | 60 | 5 | 9 | 0 | 1172.15 |
| 35 | 1 | 70 | 60 | 9 | 0 | 1172.3 |
| 36 | 0 | 60 | 5 | 6 | 0 | 1535 |
| 37 | 0 | 60 | 5 | 9 | 1 | 2193 |
| 38 | 1 | 80 | 5 | 9 | 0 | 4153.547 |
